# Structured priors in human forecasting

**DOI:** 10.1101/285668

**Authors:** Francisco Quiroga, Eric Schulz, Maarten Speekenbrink, Nigel Harvey

## Abstract

Forecasting is an increasingly important part of our daily lives. Many studies on how people produce forecasts frame their behavior as prone to systematic errors. Based on recent evidence on how people learn about functions, we propose that participants’ forecasts are not irrational but rather driven by structured priors, i.e. situationally induced expectations of structure derived from experience with the real world. To test this, we extract participants’ priors over various contexts using a free-form forecasting paradigm. Instead of exhibiting systematic biases, our results show that participants’ priors match well with structure found in real-world data. Moreover, given the same data set, structured priors induce predictably different posterior forecasts depending on the evoked situational context.

## Introduction

Forecasting plays a crucial role in our daily lives. We decide whether to take an umbrella to London based on a weather forecast. We forecast our commute time to determine when to leave home. Forecasting is also vital in the business world. For instance, sales forecasts aid companies to manage their inventories, and event production companies predict attendance numbers to select a suitable venue. Weller and Crone (2012) surveyed 173 companies from different sectors and found that 44% of their forecasts were based on human judgments aided by a statistical baseline. The rest of the forecasts were similarly divided into approaches that used pure statistics (29%) or pure human judgment (26%)^1^.

Psychological research on *judgmental forecasting* has advanced our understanding of different effects that arise when individuals produce forecasts from time series data (Lawrence, Goodwin, O’Connor, & Önkal, 2006). Reimers and Harvey (2011) highlight four well-established forecasting effects: (i) trend damping: participants underestimate linear trends, (ii) optimism: forecasts for untrended series are too high and trend damping is larger for downward facing trends, (iii) noise adding: participants add noise to their forecasts, and (iv) overestimation of autocorrelation. Frequently, these effects are also referred to as *judgmental forecasting biases*, under the assumption that participants’ forecasts are either suboptimal or —at the least— require the aid of decision support systems (see Armstrong & Collopy, 1998). However, other interpretations of these effects are also conceivable (cf., Harvey & Reimers, 2013).

This paper aims to further assess where systematic patterns in human forecasting come from and whether they are systematically biased or the results of an adaptation to the real world. We base our account on the assumption that phenomena observed in human forecasts mirror people’s experiences of patterns found in the real world. We show that participants’ forecasts are driven by *structured priors* that track naturally occurring patterns in their environments and that these priors can account for varying behavior in different forecasting scenarios.

### A model of structured priors

Our approach to assessing structured priors is inspired by Griffiths and Tenenbaum’s (2006) work on people’s priors over durations and magnitudes. By asking people to produce forecasts for common phenomena such as a person’s predicted lifespan or the number of lines in a poem, Griffiths & Tenenbaum extracted participants’ priors over different scenarios and found that the extracted forms matched, on average, those of a full-information Bayesian model.

We adopt a similar method, but instead of extracting priors over distributional shapes, our goal is to extract priors over functional shapes. More specifically, we will use a model based on recent evidence showing that human function learning can be well-explained by Gaussian Process regression (Griffiths, Lucas, Williams, & Kalish, 2009; Lucas, Griffiths, Williams, & Kalish, 2015; Schulz, Tenenbaum, Duvenaud, Speekenbrink, & Gershman, 2017). It is interesting to note that the difference between function learning and judgmental forecasting turns out to be rather small. Whereas function learning is frequently assessed sequentially, i.e., presenting inputs and asking for predictions of the output one at a time, in judgmental forecasting generally the entire dataset is revealed at once and subjects are subsequently asked to produce forecasts for future time points. Nevertheless, many of the effects found in judgmental forecasting translate to findings in the domain of function learning. For example, when DeLosh, Busemeyer, and McDaniel (1997) trained participants to learn an underlying quadratic function, they found that their average predictions fell between the true function and straight lines fitted to the closest training points. This behavior is similar to a form of (polynomial) trend damping as well as an overestimation of auto-correlation (Eggleton, 1982).

Recently, Schulz, Tenenbaum, et al. (2017) showed that the way in which participants learn complex functions can be explained by the use of compositional kernels formed by combining simpler structural components (see also Schulz, Tenenbaum, Duvenaud, Speekenbrink, & Gershman, 2016). We expand on these findings by assessing participants’ structured priors in a both a “free form” and a traditional forecasting paradigm.

### Formal description

We use Gaussian Process (GP) regression as a Bayesian framework to assess participants’ forecasts (see Schulz, Speekenbrink, & Krause, 2017, for a tutorial on Gaussian Process regression). Formally, a GP is a collection of random variables, any finite subset of which are jointly Gaussian-distributed (Rasmussen & Williams, 2006). A GP can be expressed as a distribution over functions. Let *f*: 𝒳 → ℝ denote a function over input space 𝒳 that maps to real-valued scalar outputs. These functions are assumed to be random draws from a GP:

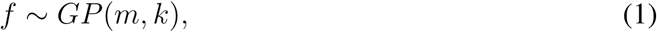

where *m* is a mean function specifying the expected output of the function for each input **x**, and *k* is a kernel (or covariance) function specifying the covariance between outputs at inputs **x** and **x**′:

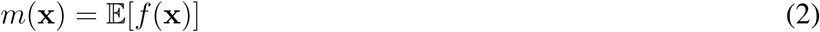

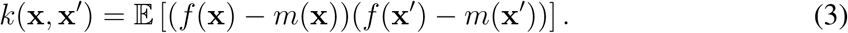

Intuitively, the kernel encodes an inductive bias about the expected shape of the underlying function (Schulz, Tenenbaum, Reshef, Speekenbrink, & Gershman, 2015). We will use the form of this kernel to assess structured priors in human forecasting. To simplify exposition, we follow standard convention in assuming a prior mean of *m*(**x**) = **0**.

Conditional on observed data 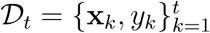, where *y*_*k*_ ∼ 𝒩 (*f* (**x**_*k*_), *σ*^2^) is a draw from the latent function with added noise, the posterior predictive distribution for a new input **x**_*_, is Gaussian *p*(**x**_*_|𝒟_*t*_) = *N* (*m*_*t*_(**x**_*_), *v*_*t*_(**x**_*_)) with mean and variance given by:

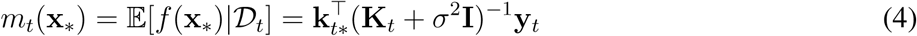

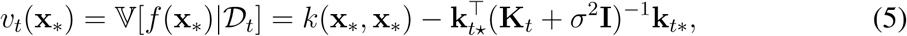

where **y**_*t*_ = [*y*_1_, …, *y*_*t*_]^⊤^, **K**_*t*_ is the *t × t* matrix of covariances evaluated at each pair of observed inputs, and **k**_*t*\*_ = [*k*(**x**_1_, **x**_*_), …, *k*(**x**_*t*_, **x**_*_)] is the covariance between each observed input and the new input **x**_*_.

Mathematically, it is possible to create richly structured and interpretable kernels by combining different base components. For example, by summing kernels, we can model the data as a sum of independent functions. Inspired by previous research on functional rule learning (Brehmer, 1974) and following recent compositional accounts of function learning (Schulz, Tenenbaum, et al., 2017), we will use three base kernels which can be combined by addition and multiplication to produce a rich set of compositional kernels. The base kernels are the linear kernel, *k*(***x, x***^**′**^) = (***x*** − *θ*_1_)(***x****′* − *θ*_1_), the periodic kernel, 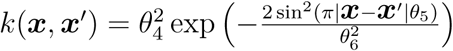, and the radial basis function kernel, 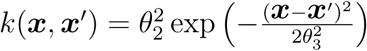. For tractability reasons, we will fix the maximum number of combined kernels to three and will not allow for repeated kernels (following Schulz et al., 2016). The complete set of resulting kernels is shown in Table 1 alongside their natural language explanations.

**Table 1.**
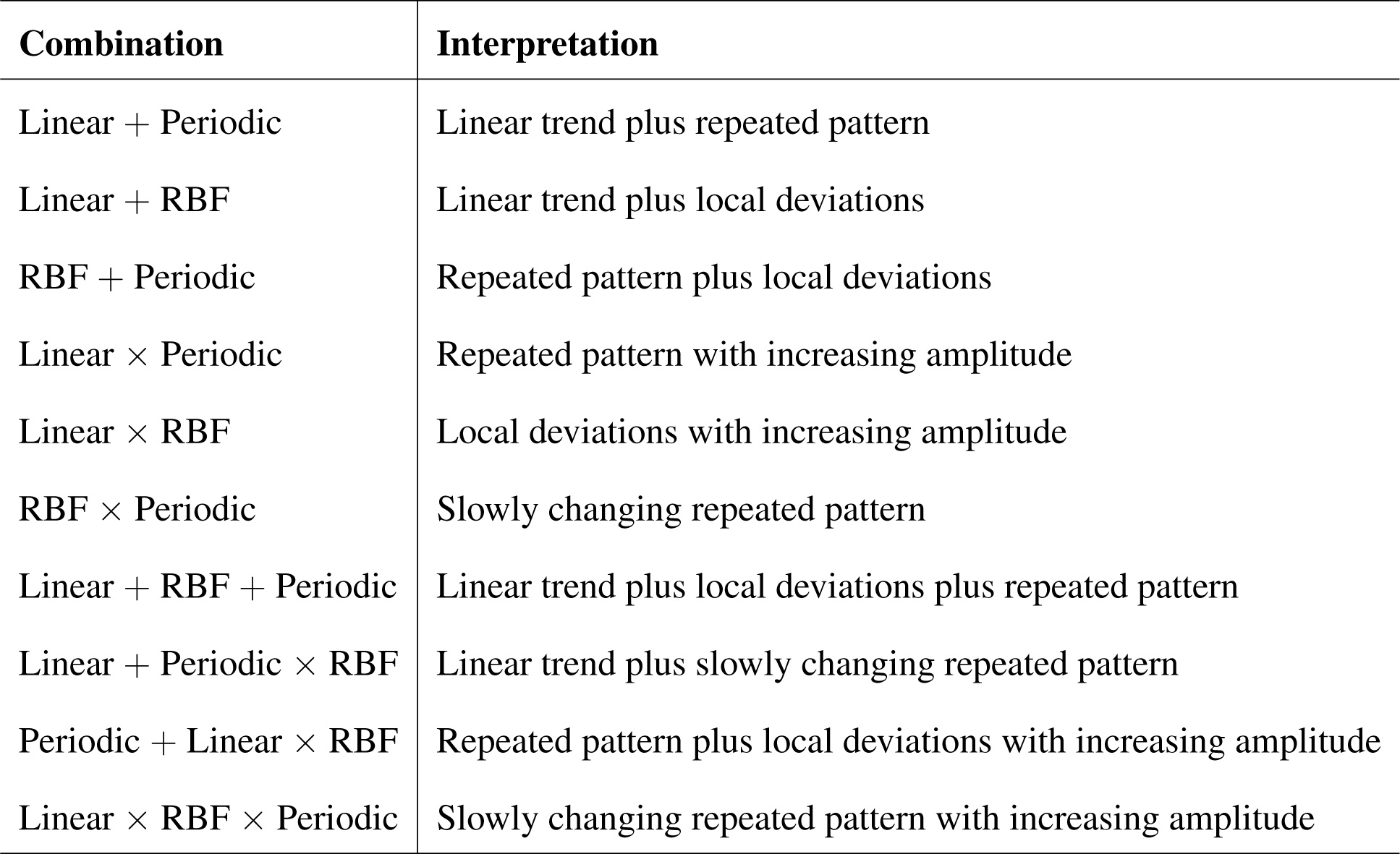
Kernel combinations in the grammar of structured functions and their interpretations.

## Experiment: Assessing prior and posterior forecasts

The present study was approved by University College London’s research ethics committee CPB/2010/013.

### Participants

121 participants (55 female) with a mean age of 34.4 (SD = 9.62) were recruited from Prolific Academic and received £1 for their participation. The experiment took an average of 14 minutes to complete.

### Set-up

Participants were asked to produce forecasts for six different scenarios. This was done by placing points over empty plots where the x-axis covered a four-year period by clicking on the plotting area. These points were automatically connected by a single curve that was computed using the centripetal Catmull-Rom method, which has the feature of ensuring that the curve passes through all the given points (Yuksel, Schaefer, & Keyser, 2009). Participants could remove points by clicking on them or by pressing a “undo” button. To avoid a curve from describing a non-injective function (i.e., having more than one value of *f* (*x*) for a single *x*), participants were asked to place their points at least one month away from each other. At the same time, participants were asked to ensure that their plots had at least one point in the first month and one in the last month (see also Figure 1). There were four conditions and six forecasting scenarios in total. Every participant encountered a balanced combination of the conditions and scenarios, totaling a number of 12 trials overall.

**Figure 1.**
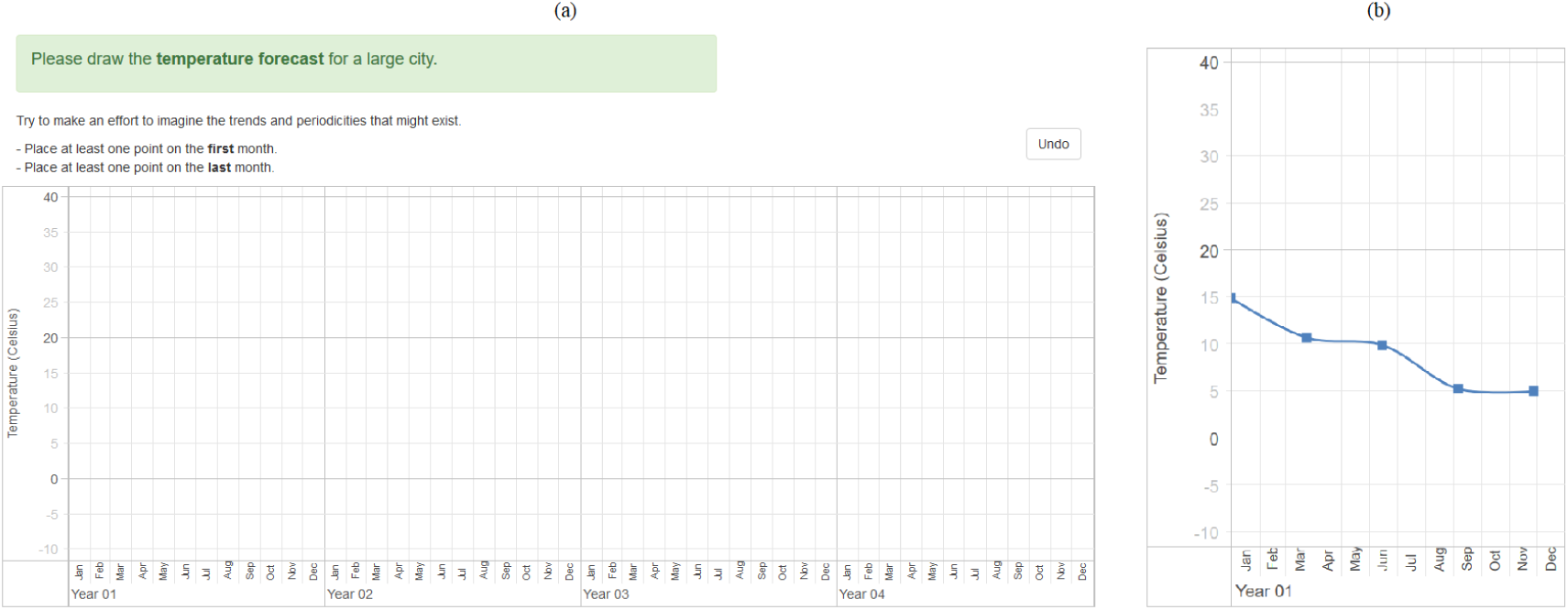
Screenshots of the experiment’s visual interface. **A**: Prior condition for the Temperature scenario. Participants could place as many points as they liked by clicking on the plotting area. **B**: Posterior condition with negative trend. Years 2 to 4 have been cut off the image but were presented to participants to enter their forecasts.

### Materials and procedure

#### Scenarios and conditions

The experiment consisted of four conditions (1+3): a Prior condition which was the same for each participant, and Posterior-Positive, Posterior-Stable, and Posterior-Negative conditions which were randomly sampled for each scenario and participant. There were six scenarios in total: participants had to forecast *temperature, probability of rain, sales, gym memberships, hourly wage*, and *number of Facebook friends*, all over 4 years in total. We intentionally chose six scenarios from relatively different contexts which included both frequently and less frequently experienced situations.

The plotting area on which participants drew their predictions consisted of four years on the horizontal axis. The range of the vertical axis depended on the scenario and were as follows: −10 to 40°C for temperature, 0 to 100% for rain probability, 0 to 5000 units for sales, 0 to 50 for gym members, 0 to 1000 for Facebook friends, and 0 to £50 for the hourly wage.

#### Stage I: Prior condition

During the first stage, every participant was sequentially presented with the six scenarios in random order. This stage was associated with the Prior condition only and designed to extract participants’ structured priors before any contact with experimental data. We used a free-form forecasting paradigm in which participants were shown an empty plotting area and instructions in text format (see Figure 1a). Each screen started with general instructions that the participant could show or hide at will. Additionally, each scenario was introduced with a single sentence explaining the context, for example: “Please draw the temperature forecast for a large city”.

#### Stage II: Posterior conditions

After participants had entered the predictions on the six scenarios in the Prior condition, the scenarios reappeared —in the same order— for the Posterior conditions. Participants had to produce predictions once again. However, the initial plotting areas were not empty anymore; five pre-established and inalterable data points were automatically displayed during this stage of the experiment. These points were equidistantly distributed on the first 11 months (i.e., from the first day of January to the last day of November of the first year; see Fig. 1b). The values of these points depended on the current Posterior condition and a noise group. The specific condition determined the slope of an underlying linear trend that could be either positive, negative, or stable (i.e., constant). The noise group determined the noise applied over the underlying linear trend. The occurrence was balanced so that each of the three Posterior conditions appeared twice to each participant (i.e., the posterior-positive condition appeared twice, and so did the posterior-stable and posterior-negative conditions).

The curves for two different scenarios under the same Posterior condition and noise groups were displayed to be visually equivalent; only their intercept with the y-axis differed (see Appendix B for details).

#### Behavioral results

Two participants were not included in the analysis. One consistently placed one point at the minimum value for the last month, and the other indicated browser problems.

We interpolated participants’ submitted points using the same centripetal Catmull-Rom method that was used in the experiment. We did this because the interpolation was presented to the participants in real time during the experiment. Hence it is natural to include it as part of each participant’s input (see Fig. 1). Moreover, applying the same interpolation method ensures getting the same points that participants saw, whereas other interpolation methods could produce different curves for the same dataset.

The data associated to the first 30 days (in the Prior condition) and last 30 days (in all conditions) of the four years were filtered out of the analysis to ensure that every selected day had a value for every participants’ predictions. This was because, although participants were forced to place one point in the first month and one in the last month, they could do so *in any day* of those months. Hence, one participant’s prediction might have her first value on the 1st of January while another one might have hers on the 31st of January. Lastly, we used data points in steps of five days (e.g., 31, 36, 41).

Figure 2A presents all the behavioral data collected in the Prior condition. We can see that some scenarios present strong periodicities in the Prior condition (e.g., Temperature, Rain) while others do not (e.g., Salary, Facebook friends). Moreover, participants visibly generated different time series for different scenarios, thereby indicating that they might have different prior expectations about the shape of the time series depending on the scenario.

**Figure 2.**
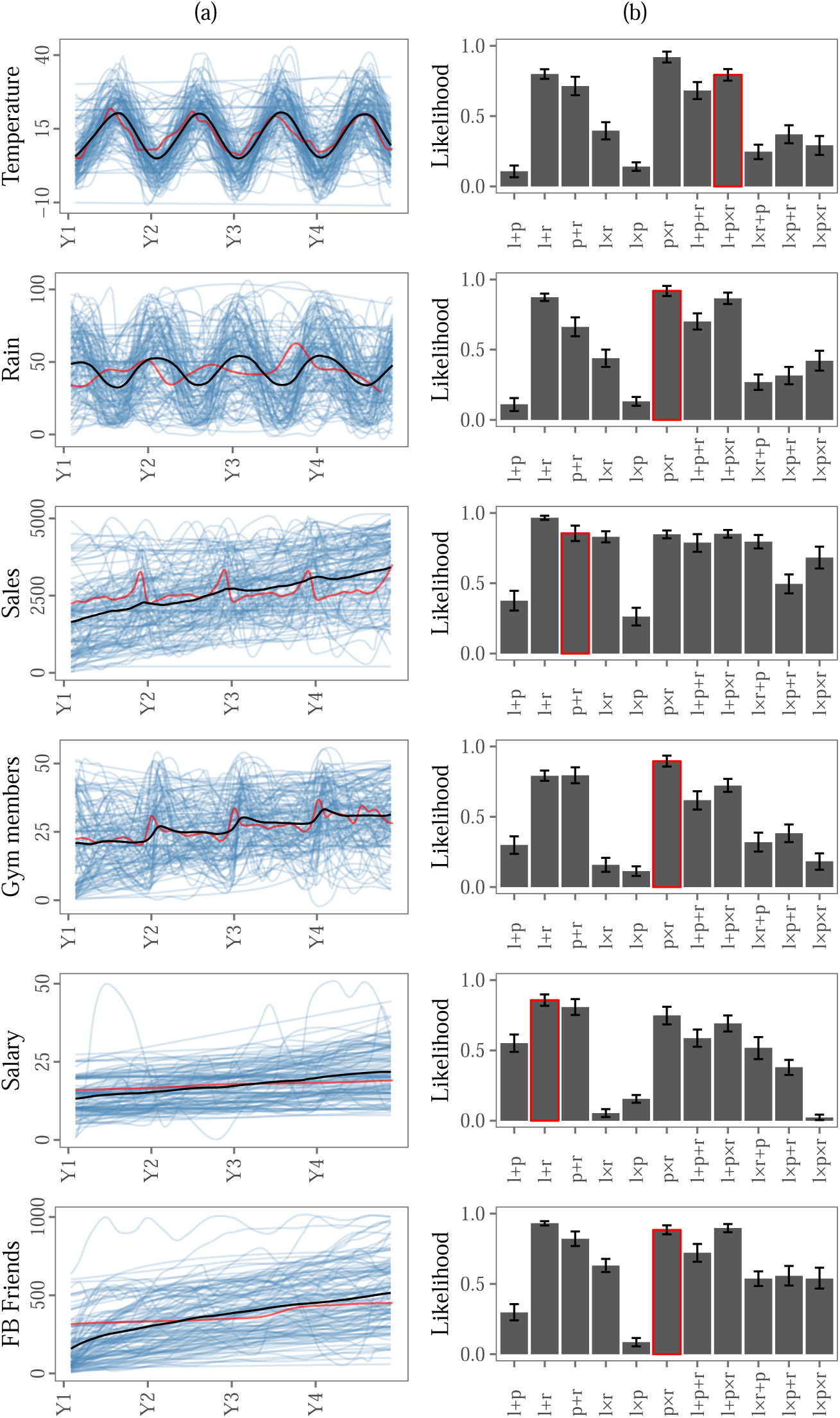
**A**: Results of Stage 1. Blue curves represent each participant’s input while the black curves represent the mean value averaged over participants. Red curves represent real world data taken from related domains, mean-centered by using the empirical means. **B**: Standardized prior likelihoods over compositions, averaged over participants. Error bars represent the standardized 95% confidence interval of the means. Red borders indicate the best fitting composition for the real world data.

### Extracting structured priors

We fitted Gaussian Process (GP) regressions, with each of the eleven possible compositional kernels, to every forecast produced by each participant in the six Prior condition scenarios using the GPFlow software (Matthews et al., 2017).^2^ In cases where the GP conditioning procedure failed to optimize (often due to problems with the Cholesky decomposition), a white noise kernel was added to the composition. If the GP failed to optimize again, it was flagged for removal. We also removed those log marginal likelihoods that were five standard deviations or more away from the mean of each scenario-kernel combination. Overall, 5.02% of the fitted GPs were excluded using this procedure.

For each participant’s forecasts (i.e., for each participant-scenario-condition combination), we standardized the likelihoods produced by each of the GPs’ kernel compositions to be in the range from 0 to 1 to approximate the posterior probabilities assuming equal prior probability for each kernel combination. Afterwards, we averaged the results, per kernel and scenario, over participants. We executed this process to obtain an approximate compositional prior over the different kernels, as currently there is no reliable way to reach actual distributions over combined kernels (but see Janz, Paige, Rainforth, van de Meent, & Wood, 2016). The resulting approximated prior distributions are shown in Figure 2B.

We can see that the different scenarios lead to diverse characteristic forecasting patterns and distinct best fitting kernels as measured by the standardized likelihood. In the temperature, rain, and gym members scenarios, participants’ prior forecasts were best modeled by a multiplicative combination of a radial basis function and a periodic kernel indicating that they assume the data to have a periodicity that is not always perfectly repeating. In the sales, gym membership, and Facebook friends scenarios, participants’ priors were best captured by an additive combination of a linear and a radial basis function kernel. This indicates that people might expect this data to have a trend, yet that this trend does not persist indefinitely but slows down over time.

### Human priors track real-world structure

We found that the kernels that best described the data in the Prior condition depended on the scenario. To understand whether this might be due to a de facto match between a scenario’s actual structure and participants’ priors, we gathered real-world data and processed it.

We used four years of real-world data for each scenario, using proxies when direct data was unavailable. In particular, the temperature data was the daily temperature in London from 2013 to 2016. The rain data was the daily rainfall data in the city of London for the same period. We summarized the rain data by trimesters to estimate the probability of rain over time. The sales data was obtained from a report by the Office for National Statistics of the United Kingdom which includes all retail sales (excluding fuels). This time series was monthly for the years 2013 to 2016. The gym membership data was generated by exporting how many times people searched for the term “gym membership” on Google, from 2013 to 2016. For the salary data, we used the median annual earnings, as reported by The Hamilton Project^3^, of adults ranging from 25 to 28 years old (as per the question participants had to answer; see Appendix A). Finally, the Facebook friends data was exported from one of the author’s actual Facebook profile, starting from when he was 25 years old. We processed every real-world dataset as if it was participants’ data; the Catmull-Rom interpolation was calculated to produce daily values, and the different Gaussian Process kernels were fitted to the data.^4^

Figure 2 shows red rectangles around the kernels that best described the collected real-world data. The optimal kernel composition coincided for the participants and the real-world data in the Rain, Gym members, and Salary scenarios. In the Temperature scenario, the best fitting kernel for the real-world data was *l* + *p × r*, which has a linear component that the optimal kernel composition for the participants’ data did not have. The Sales and Facebook friends scenarios best fitting kernels in the real-world data were *p* + *r* and *p × r*, respectively. In the case of Facebook friends, although the participants’ best fitting kernel was *l* + *r*, the standardized likelihood for *p × r* was not statistically different.

Next, we assessed the rank correlation between each participant’s prior distribution over compositions and the distributions derived from the real-world data for each scenario. The average rank-correlation per scenario was *ρ* = 0.61 (*t*(118) = 32.26, *p <* 0.001) for the Temperature data, *ρ* = 0.58 (*t*(118) = 24.63, *p <* 0.001) for the Rain data, *ρ* = 0.48 (*t*(118) = 20.23, *p <* 0.001) for the Sales data, *ρ* = 0.16 (*t*(118) = 6.41, *p <* 0.001) for the Gym members data, *ρ* = 0.61 (*t*(118) = 37.3, *p <* 0.001) for the Salary data, *ρ* = 0.62 (*t*(118) = 33.62, *p <* 0.001) for the Facebook friends data. Thus, we found a good match between participants’ structured priors and the structured priors extracted from real-world data.

### The effect of priors on posterior forecasts

In the Posterior conditions, participants were shown 11 months of data and were asked to forecast the time series for the remaining three years. We expected participants to *push through* their ideas of how the world works from the Prior to the Posterior conditions, i.e., perform a sensible update of their structural expectations. If this was the case, then we expected to observe varying participants’ predictions for data that is visually identical but presented as stemming from different scenarios. Moreover, the better a kernel performed in the Prior condition, the better it should be able to predict participants’ forecasts in the Posterior conditions.

We fitted new GPs to the 11-month of data presented to the participants. Adhering to a Bayesian treatment, we informed these new GPs with the optimal hyper-parameters found in the respective GPs of the Prior condition. For example, the hyper-parameters that were found to maximize the fit of a GP with kernel *p* + *r* to the forecasts of Participant A in the Prior condition Temperature scenario were subsequently used as *prior values* of a new GP (with the same *p* + *r* kernel) that was fitted to the data that Participant A saw in the respective Posterior condition for the Temperature scenario. The prior distribution for each hyper-parameter was assumed to be Gaussian with a mean value equal to the optimal value found in the Prior condition, and a variance of 1. This procedure was performed for every scenario-participant-kernel combination.

Following the same data preparation method as in the Prior condition, the GPs that failed to optimize, and those with likelihoods five standard deviations or more away from the mean of each scenario-kernel combination, were removed. In total, these 1.66% of the Posterior estimations were removed this way.

We used the new GPs to produce predictions from month 12 onwards, in line with the participants’ experimental task. Afterwards, we compared these predictions to each participant’s actual forecasts and calculated the mean squared error between the models’ and participants’ predictions (see Mozer, Pashler, & Homaei, 2008, for a similar approach). Finally, we assessed the correlation between a kernel’s fitting performance in the Prior condition against its predictive performance in the Posterior condition. The former was operationalized as the standardized likelihood computed in the section above. The latter was operationalized as a rank over the error of each kernel composition.

Figure 3B shows the results of this analysis. The average correlations between a kernel’s prior likelihood and its error in capturing participants’ forecasts was *ρ* = −0.23 (*t*(118) = −6.77, *p <* 0.001) for the Temperature data, *ρ* = −0.25 (*t*(118) = −8.54, *p <* .001) for the Rain data, *ρ* = −0.1 (*t*(118) = −3.37, *p* = .001) for the Sales data, *ρ* = −0.16 (*t*(118) = −4.6, *p <* .001) for the Gym membership data, *ρ* = −0.14 (*t*(118) = −4.35, *p <* .001) for the Facebook friends data. In the case of the Salary scenario, the correlation was not significantly different from zero at *ρ* = −0.01 (*t*(118) = −0.24, *p* = .81). Thus, we found a match between participants’ structured priors and those structures’ performance when describing their Posterior condition forecasts. These results indicate that participants priors extracted from the first stage continued to influence their predictions in the second stage. At the same time, this analysis also shows that participants can adapt the structure of their forecasts in light of new evidence. As posterior forecasts should always be influenced by both prior assumptions as well as the incoming data, one would indeed expect a medium —and not a strong— match between the extracted structured priors and these structures’ performance in the Posterior condition.

**Figure 3.**
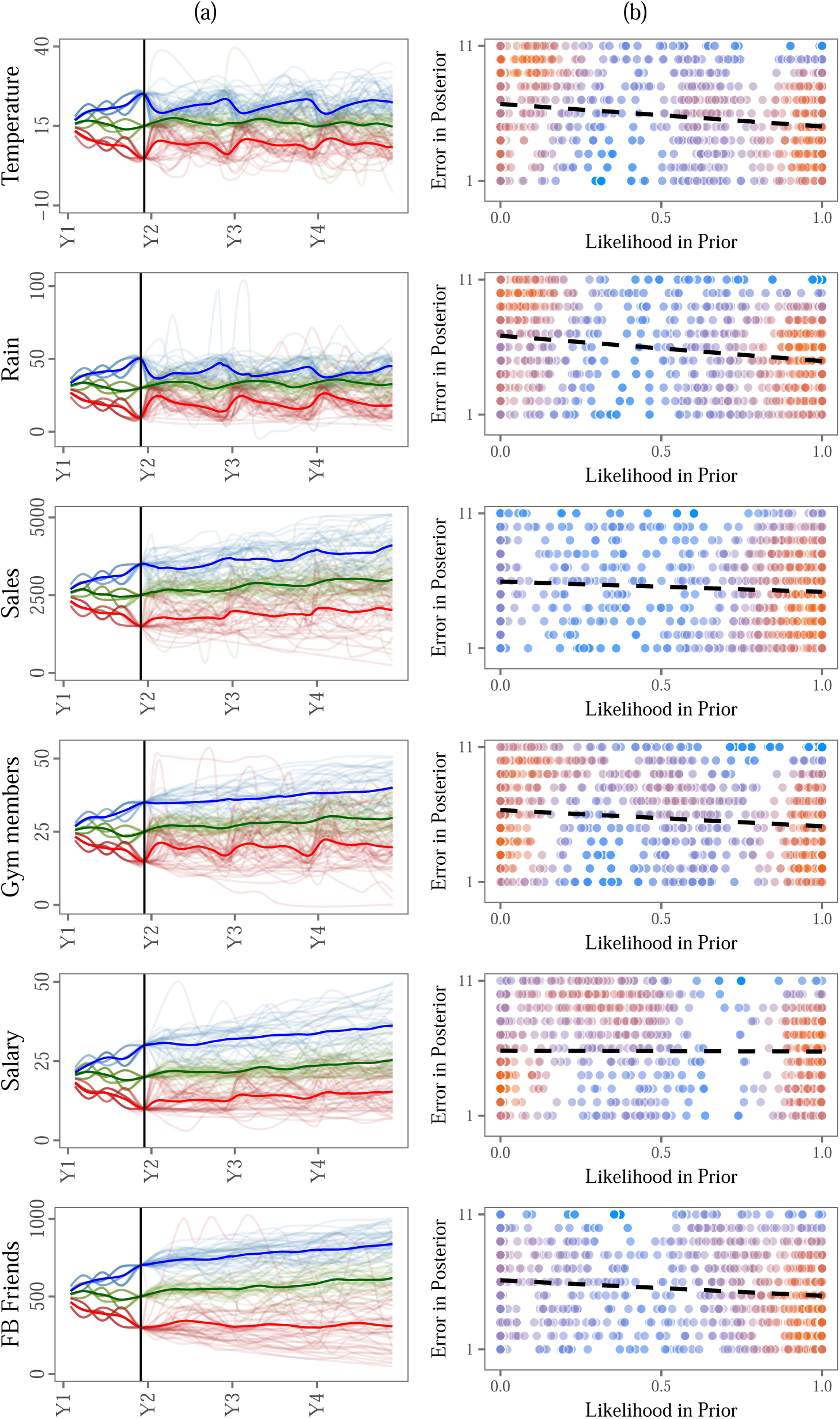
Results of Stage II. **A:** Participants’ actual forecasts for each scenario in the different conditions (blue: positive, green: stable, red: negative). **B:** Standardized likelihood in the Prior condition condition against its respective rank of error in the Posterior condition.

## Discussion and Conclusion

We proposed a novel experimental paradigm and a Bayesian structure search model to study the relationship between the forecasts that humans make before and after contact with experimental data. Our results revealed a good match between participants’ structured priors and the structure found in real-world data. This indicates that participants’ priors across a diverse set of forecasting scenarios do not reflect a manifestation of a behavioral bias, but rather result from an intelligent adaptation to their environment. Moreover, participants’ priors are strong enough to induce predictable differences in their forecasts for visually similar time series data presented in different scenarios.

We are not the first to find evidence that people’s forecasts might be representative of their environment. In fact, in the context of judgmental forecasting, Harvey and Reimers (2013) showed that the trend damping effect can be a result of an adaptation to the environment. This evidence opposed two other hypotheses that indicated an anchoring bias or an experimental interference. We believe that our results strengthen the idea of adaptation over bias.

### Limitations

Currently, our experimental setup suffers from at least three shortcomings: (i) the set of presented scenarios could be larger, (ii) the ranges shown in the plots were pre-established and fixed, and (iii) the information in the Posterior condition was presented statically. First, we only assessed participants’ priors across six different scenarios. Although we selected them from a relatively broad range of topics, humans are capable of producing forecasts in vastly more diverse and richer settings than those assessed here. Second, we acknowledge that, despite our best efforts, pre-established ranges will always exert an influence on participants’ forecasts. Finally, we approximated the underlying distributions over kernels by standardizing the marginal likelihood produced by each kernel composition for every participant. This rough approximation was chosen as current implementations providing fully Bayesian treatments of estimating posteriors over kernels converged unreliably and very slowly in our datasets. Thus, we intend to reassess our current data by calculating the true posterior over structures as soon as the required software to do so becomes available.

### Future Work

In future work, we intend to further broaden the scope of our proposal by assessing participants’ behavior in larger and more diverse sets of forecasting scenarios while also adding more kernels to our grammar. One interesting question for follow-up studies could be to contrast unfamiliar with familiar scenarios thereby assessing how priors develop as participants gain experience with a scenario.

Additionally, we believe that the effect of presentation formats on priors in time series data should also be assessed. For example, Kusev, van Schaik, Tsaneva-Atanasova, Juliusson, and Chater (2018) showed that presenting time series data sequentially instead of revealing all of the points at once improved the accuracy of participants’ forecasts but did not have an effect on their estimations of the mean.

Finally, inspired by the results of Harvey and Reimers (2013), and considering that the RBF kernel has the characteristic of reverting the predictions to the mean, we hypothesize that our paradigm could eventually explain trend damping. In fact, we tested this hypothesis with our current setup yet found no conclusive evidence. Nevertheless, we believe that this is a natural step forward.

## Conclusion

Using a Bayesian model of structure search, we have found that participants’ priors in different forecasting scenarios are structural by nature, are well-aligned with the underlying structure of the environment and lead to predictable differences in forecasts for the same data when labeled as coming from different scenarios. These results enrich our understanding of how people generate forecasts by using both prior structural expectations and the encountered data.

## Acknowledgements

FQ was supported by the *Becas Chile* program of the National Commission of Scientific and Technological Research (CONICYT). ES is supported by the Harvard Data Science Initiative.

## Appendix A

### Supplementary Materials

The following are the questions that participants encountered during the first stage of the experiment, depending on the scenario.

- Please draw the **temperature forecast** for a large city.
- Please draw the **sales forecast** for a large company.
- Please draw a graph showing the **number of total Facebook friends** of a 25 year old male.
- Please draw the **probability of a rainy day** for a large city.
- Please draw the **number of total gym members** of a small gym.
- Please draw the **hourly wage** of a 25 year old male.

The following sentence was added to the second stage (i.e., the Posterior conditions) of the experiment: *given the information for the first year*.

### Appendix B

Initial values in the Posterior conditions

#### Y-intercept value

The Y-intercept value for Stage II of each scenario were chosen as *reasonable* values by the experimenters. These values were 15 for temperature, 30 for rain, 2500 for sales, 25 for gym member, 500 for Facebook friends, 20 for hourly wage.

#### Noise values

There were three noise groups of five elements each (see Table B1). Every element was sampled from a Uniform distribution with minimum value 0 and maximum value 1. These random numbers were generated in Python 3.5.2 by `random.random()`, from the `random.py` library. Each noise group was then normalized to have a mean of zero and a standard deviation of 1. Finally, the three center elements of each group were added to the linear trend. The details of the last two steps are explained again further in this Appendix section.

**Table B1.** Noise groups used in the creation of the Posterior conditions’ initial values. Values have been rounded to two decimals; complete multi-decimal values can be found on GitHub.

| Noise group | Values of the noise group |
| --- | --- |
| 0 | 0.76, 0.59, 0.80, 0.36, 0.43 |
| 1 | 0.45, 0.77, 0.09, 0.34, 0.81 |
| 2 | 0.62, 0.04, 0.06, 0.04, 0.17 |

#### Initial values’ calculation

The Y-intercept values, noise group, and posterior condition were integrated in such a way that when participants looked at two plots with the same posterior condition and noise group (e.g., posterior-positive and group 1, respectively), both plots would look equivalent (albeit shifted to have lower or higher values). However, as is clear by focusing on the values of the different plots in Figure B1, the curves are not quantitatively equivalent.

**Figure B1.**
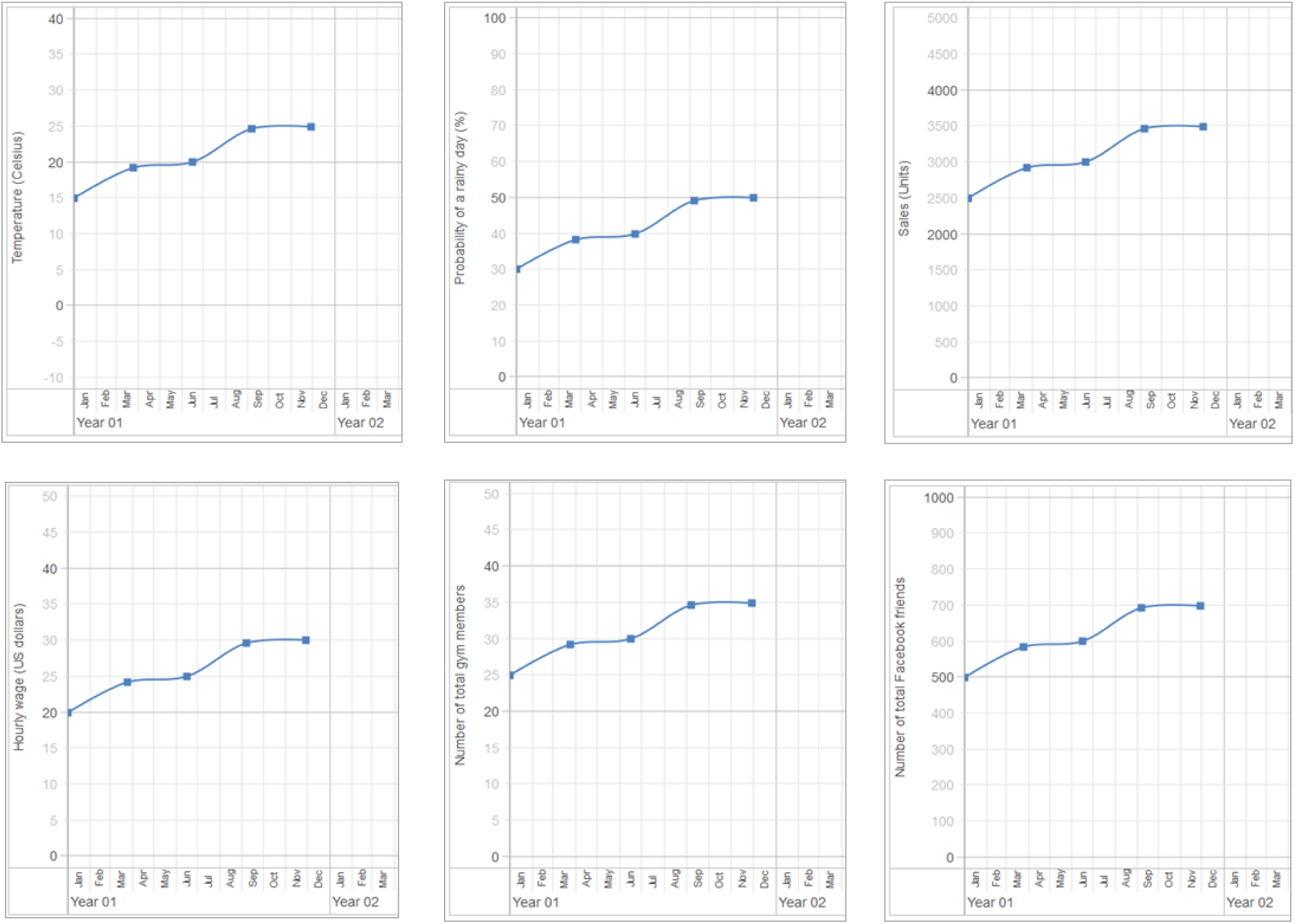
Screenshots of the six scenarios under the same posterior group and noise group. It can clearly be seen that despite each scenario having different ranges in the Y-axis, the curves *look* the same.

The five points of the first year are described by the following formula:

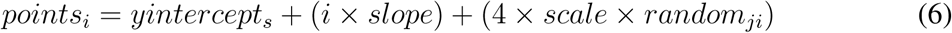

where:

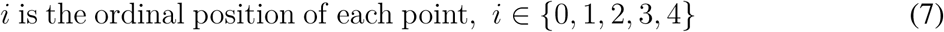

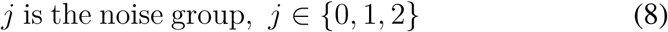

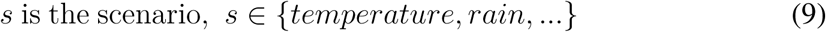

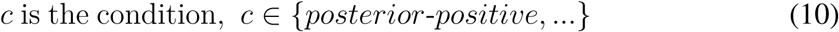

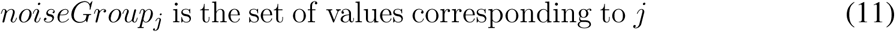

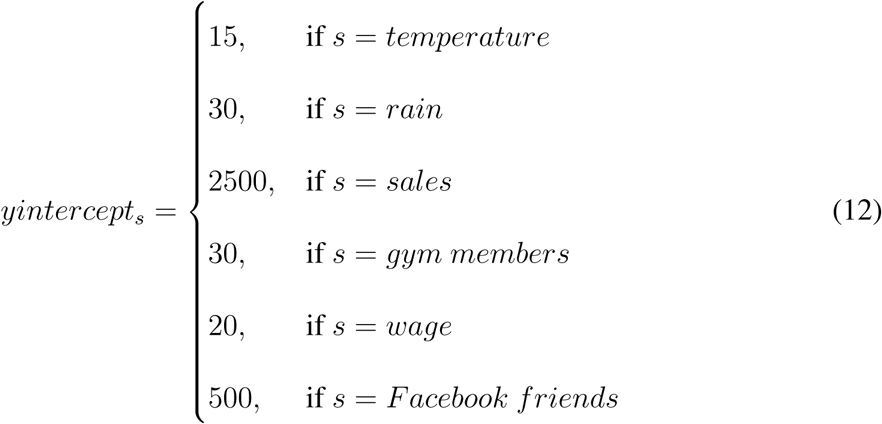

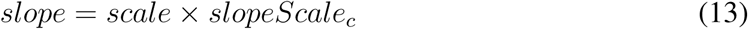

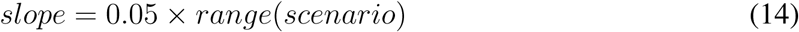

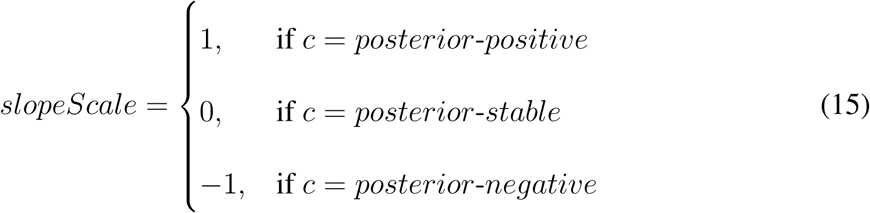

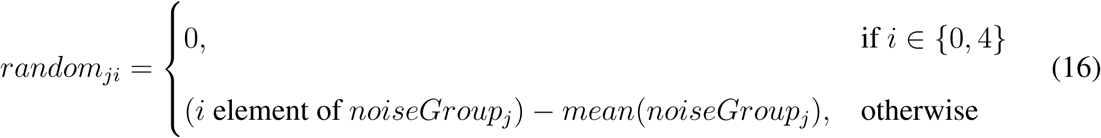

## Footnotes

1 The percentages in the original study do not add up to 100%, presumably due to rounding.

2 The optimization time of each GP took approximately 1 minute on average on a personal computer (Intel(R) Core(TM) i5-6200U at 2.30 GHz, 8GB of RAM, running Python 3.6.1). This lead to an overall runtime of 11 hours for the 714 combinations.

3 http://www.hamiltonproject.org

4 All data can be found online: https://git.io/vA86Q

